# *TelNet* - a database for human and yeast genes involved in telomere maintenance

**DOI:** 10.1101/130153

**Authors:** Delia M. Braun, Inn Chung, Nick Kepper, Katharina I. Deeg, Karsten Rippe

## Abstract

The ends of linear chromosomes, the telomeres, comprise repetitive DNA sequences that are protected by the shelterin protein complex. Cancer cells need to extend these telomere repeats for their unlimited proliferation, either by reactivating the reverse transcriptase telomerase or by using the alternative lengthening of telomeres (ALT) pathway. The different telomere maintenance (TM) mechanisms appear to involve hundreds of proteins but their telomere repeat length related activities are only partly understood. Currently, a database that integrates information on TM relevant genes is missing. To provide a reference for studies that dissect TM features, we here introduce the *TelNet* database at http://www.cancertelsys.org/telnet/. It offers a comprehensive compilation of more than 2,000 human and over 1,100 yeast genes linked to telomere maintenance. These genes were annotated in terms of TM mechanism, associated specific functions and orthologous genes, a TM significance score and information from peer-reviewed literature. This TM information can be retrieved via different search and view modes and evaluated for a set of genes on a statistics page. With these features TelNet can be integrated into the annotation of genes identified from bioinformatics analysis pipelines to determine possible connections with TM networks as illustrated by an exemplary application. We anticipate that *TelNet* will be a helpful resource for researchers that study TM processes.

## Background

Telomeres, the ends of linear chromosomes, consist of repetitive DNA sequences bound by the shelterin protein complex [1, 2]. This protein assembly protects the DNA ends from degradation and accidental recognition as DNA double-strand breaks [3-5]. The progressive shortening of the telomere repeat sequences that accompanies normal replication limits the number of cell divisions. Thus, it needs to be circumvented by cancer cells for unlimited proliferation. This is accomplished by activation of a telomere maintenance (TM) mechanism. It involves either the reactivation of the reverse transcriptase telomerase normally repressed in somatic cells via different mechanisms [6-9], or activation of the alternative lengthening of telomeres (ALT) pathway [10-13]. ALT activity in human cancer cells occurs via DNA repair and recombination pathways but details on the mechanism remain elusive. Thus, TM is a complex process that involves proteins that are located as part of the shelterin complex at telomere repeats [14, 15] or in close proximity [16, 17]. Factors that regulate transcription and functional activity of telomerase are relevant [18, 19] as well as features of the ALT pathway like PML (promyelocytic leukemia) nuclear bodies along with telomere repeats that are associated with a variety of proteins and referred to as APBs (ALT-associated PML nuclear bodies) [20-23]. Furthermore, investigations of deregulating effects upon telomere shortening have related a number of proteins to telomere crisis [24].

A well-studied model organism for telomere biology is the budding yeast *Saccharomyces cerevisiae* [25]. Several independent deletion screens with subsequent direct measurements of telomere length have identified a comprehensive list of yeast genes involved in telomere length regulation [26-28]. Since telomere structure and function are highly conserved between organisms, mammalian homologues exist for most of the genes identified in the various yeast screens. Thus, it can be informative to relate TM phenotypes found in yeast to human homologues [29]. In *S. cerevisiae*, telomerase is constitutively active and its deletion leads to cellular senescence [30]. Survivor cells that overcome cellular senescence in the absence of telomerase use a mechanism based on homologous recombination for telomere elongation [31]. Interestingly, similar to ALT in human cells, so-called type II survivors are characterized by heterogeneous telomere lengths [32, 33].

The following existing databases provide some telomere-relevant information: *TeloPIN* (Telomeric Proteins Interaction Network, http://songyanglab.sysu.edu.cn/telopin/index.php) is a collection of interaction data in human and mouse cells from available literature and GEO (gene expression omnibus) database [34]. It includes interactions with shelterin roteins and protein interactions of the non-coding telomere repeat-containing RNA TERRA). It also provides information on the methods used to identify the respective interaction. The *Telomerase database* (http://telomerase.asu.edu/overview.html) is a web-based tool for the study of structure, function, and evolution of the telomerase ribonucleoprotein [35]. It is a comprehensive compilation of information on the telomerase enzyme and its DNA substrate. The previously published *TeCK database* (http://www.bioinfosastra.com/services/teck/index.html) is a collection of telomeric and centromeric sequences as well as telomerase, centromere and kinetochore binding proteins [36] but appears to be no longer available. Finally, *MiCroKiTS* (Midbody, Centrosome, Kinetochore, Telomere and Spindle; http://microkit.biocuckoo.org) provides information on the cellular localization of proteins relevant for cell cycle progression and also includes telomere proteins [37].

All the above-mentioned databases cover telomere related information but lack an annotation of genes with respect to the TM mechanism. Accordingly, we here introduce a compilation of information on telomere maintenance relevant genes via the *TelNet* database. *TelNet* currently comprises more than 2,000 human, and over 1,100 yeast genes that are involved in TM pathways. The annotation of these genes includes the classification of telomere maintenance mechanisms along with a significance score as well as TM specific functions and homology assignments between different organisms. Furthermore, links to the respective literature sources are given. Thus, *TelNet* provides an integrative platform for dissecting TM networks and elucidating the alternative lengthening of telomeres pathway.

## Construction and content

### Implementation

The *TelNet* database was constructed using the Filemaker Pro software version 13. It is accessible at http://www.cancertelsys.org/telnet and is distributed via Filemaker server version 15 via its webdirect module. The *TelNet* webpage provides general information about *TelNet* as well as instructions on how to use it. Links to other databases and contact information are provided as well.

### Data source

To compile an initial set of TM relevant genes, we selected screening studies on genes or proteins that play a role in telomere biology (Figure 1, Table 1) that included the following: (i) Proteins that bound to a telomere probe in an ALT- and a telomerase-positive cell line [14], (ii) proteins from the analysis of telomeric chromatin of telomerase-positive cells [15], (iii) proteins in close proximity to shelterin components [16, 17], (iv) proteins that affected ALT-associated PML nuclear bodies [23, 38], (v) deregulated proteins linked to telomere shortening [24], (vi) genes identified from telomerase activity signatures derived from gene expression data [39], (vii) telomerase regulators identified from a kinase screen or reviewed recently [18, 19] and, (viii) a gene set with potential relevance to telomeres and the ALT pathway [40]. In addition, more than 1,100 budding yeast genes were included in *TelNet*. These genes were identified from the following sources: (i) Deletion screens that identifying telomere length associated genes [26-28], (ii) post-senescent survivor screening after telomerase knockout [41] and, (iii) transcription factors of telomerase [42].

**Figure 1.**
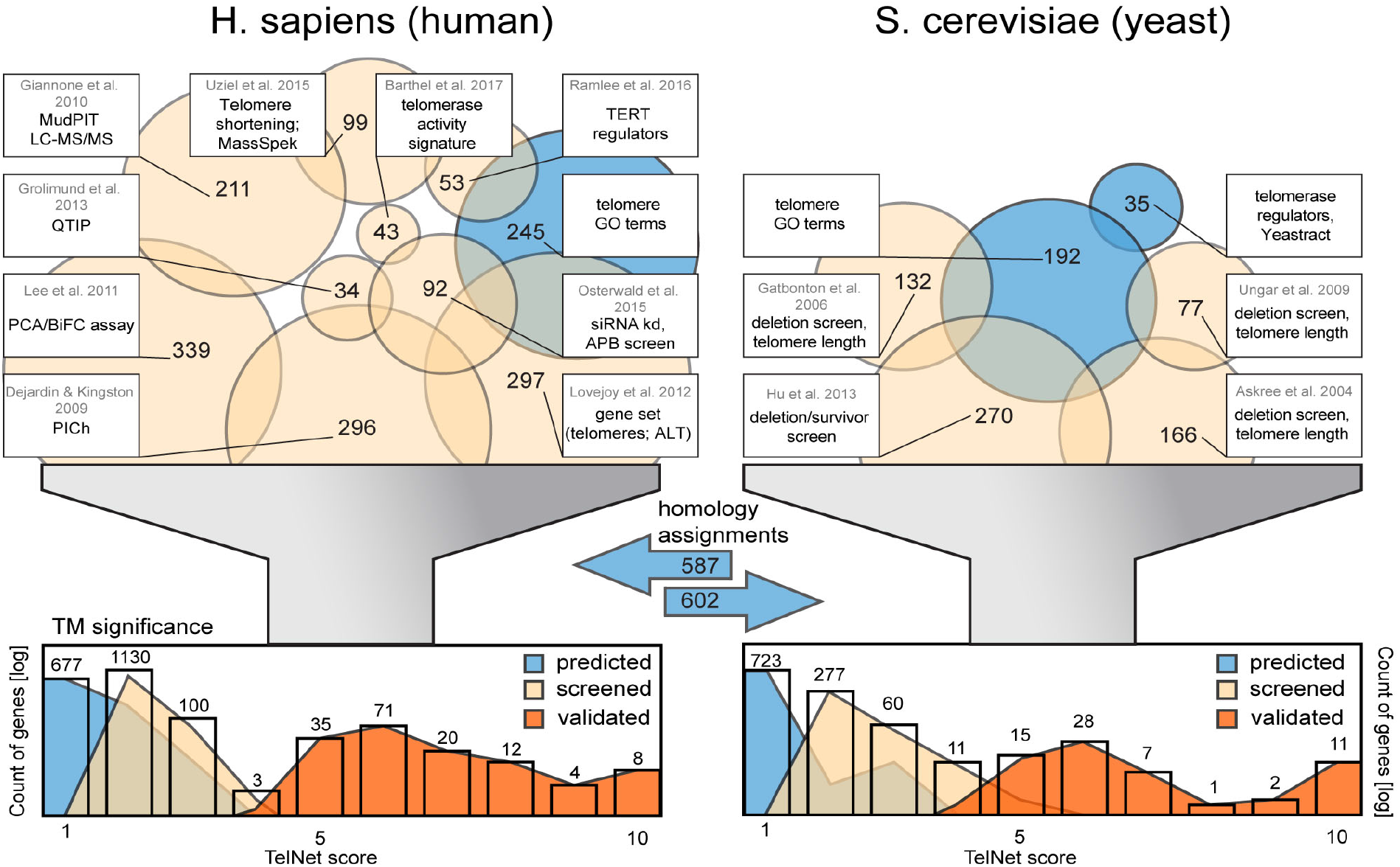
Data sources of TM genes for *TelNet*. Selected screening studies and other references that serve as sources for TM genes are indicated. In total over 2,000 human genes and more than 1,100 budding yeast genes were included in the *TelNet*. Histograms of the *TelNet* scores are displayed for the complete gene sets per species, colored by TM significance. Color scheme: blue, predicted TM genes; orange, genes from screening studies; red, validated genes.

**Table 1.**
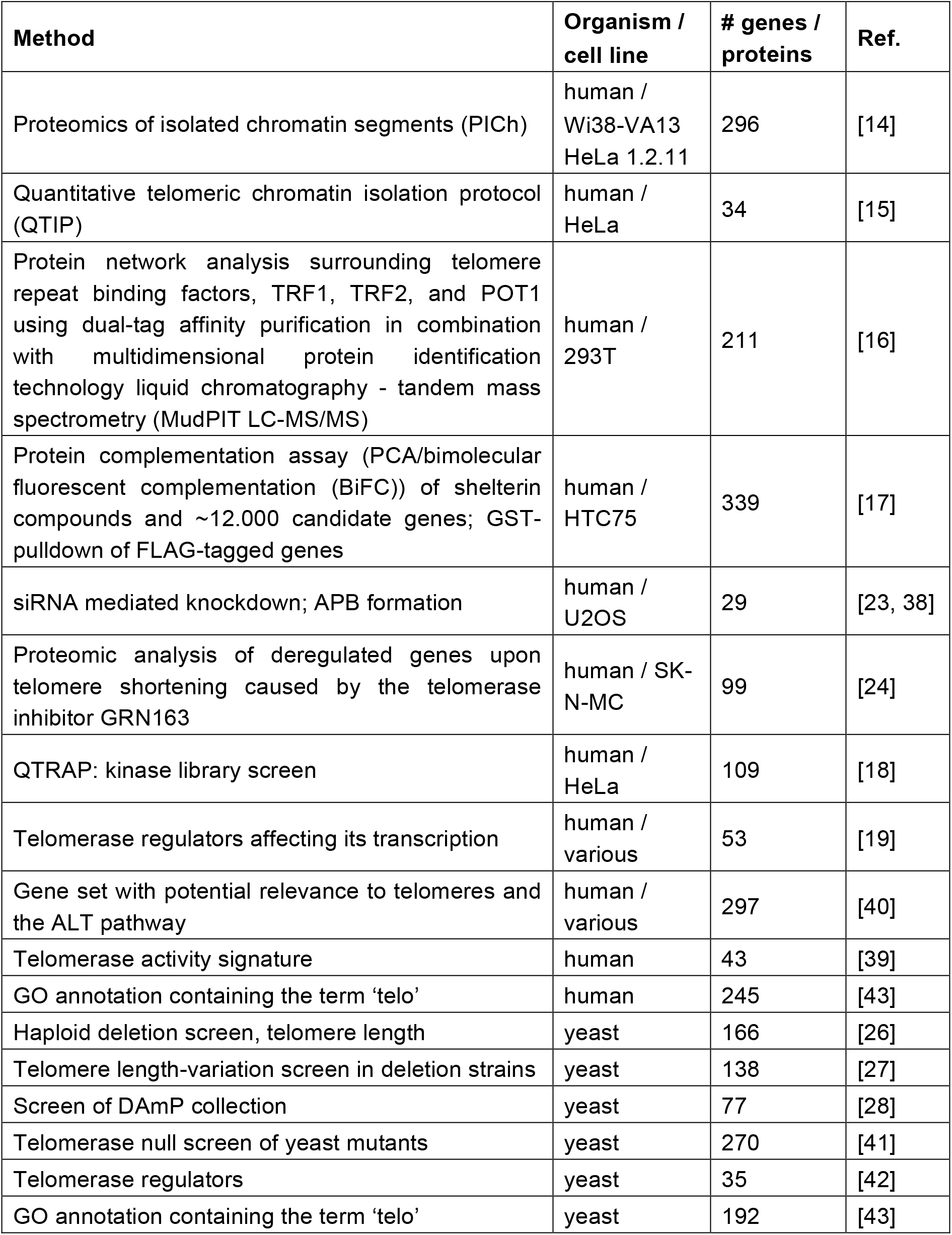
Screening studies included in *TelNet* for identification of TM genes. Selected screening studies and other sources with information on TM genes are shown. The method used for the respective investigation as well as the organism and if available the human cell line is indicated. Also, the number of genes, added from the given literature source is listed.

To differentiate the relevance of a gene or corresponding protein we introduced three categories of TM significance: ‘predicted’, ‘screened’ and ‘validated’. The factors collected from the above-mentioned screening or review sources were classified as ‘screened’. Genes with a suggested role in telomere maintenance but lacking experimentally validation were assigned with the TM significance ‘predicted’. Those with detailed experimental evidence for a connection to telomere maintenance were ranked as ‘validated’. Orthologues of gene’s classified as ‘screened’ or ‘validated’ were included in the *TelNet* database as ‘predicted’, if no further information was available. Moreover, all human and yeast genes with a GO annotation containing the term ‘telo’ [43] were included into the *TelNet* database. In this manner, we compiled an initial list of human and yeast genes that was further curated and annotated manually.

### General information from external databases

For a standardized nomenclature [44], the converter system from DAVID Bioinformatics Resources (https://david.ncifcrf.gov/) [45] or the BioMart tool from Ensembl [46] were used to provide gene and protein identifiers for Entrez, Hugo, Ensembl, Refseq and UniProt. To account for organism specific differences such as the lack of splicing and therefore isoforms in yeast or the absence of locus tags in human, the identifiers were selected differentially for each species. General gene information was retrieved from designated external databases and repositories, such as the National Center for Biotechnology Information (NCBI, https://www.ncbi.nlm.nih.gov) [47], HUGO Gene Nomenclature Committee (HGNC, http://www.genenames.org) [48], Ensembl (http://www.ensembl.org/index.html) [49, 50], and the Saccharomyces Genome Database (SGD, http://www.yeastgenome.org) [51]. The approved gene symbol, full name, and synonyms were taken from NCBI. UniProt [52] and Yeastmine [53] were consulted for the description of the cellular function in human and yeast, respectively, and assignment of orthologues was done with YeastMine. Based on the Gene Ontology (GO, http://www.geneontology.org) annotations [43] and in line with biocuration guidelines [54] we generated a list of cellular functions similar to curation procedures at SGD [55]. Every protein was manually annotated with the respective term that was most representative for its cellular function. In this manner, general information for every gene entry was compiled from a variety of external databases.

### Telomere maintenance annotation with literature information and scoring

Genes were further annotated with TM information from peer-reviewed literature for assigning them to functional categories (Figure 2). Up to five TM functions of an assembled list that comprises molecular functions as well as cellular processes and structures with regard to TM can be selected. A knock-out or knock-down phenotype related to TM features such as alterations in telomere length, increased or decreased ALT hallmarks, or effects on telomerase was described as free-text in the field ‘TM phenotype’. Details from the literature were summarized in the field ‘TM comment’. To quantify the significance of a given gene for TM we introduced the *TelNet* score ranging from 1 (low) to 10 (high) that is automatically calculated from information entered into the *TelNet* database (Table 2). Scoring criteria include the cellular function, number and relevance of assigned TM functions, and the amount of experimental data associated with the TM function of a given gene. Information of the proteins activity in the TM mechanism (TMM) is collected in the ‘TMM annotation’. For human genes, it is distinguished between alternative lengthening of telomeres (ALT) and telomerase regulation. Yeast genes can be annotated as type I and II survivors or in respect to telomerase regulation. Regulative activities were assigned as ‘repressing’, ‘enhancing’, or ‘conflicting’. The latter refers to cases where literature information was inconsistent or if genes were mentioned in the context of ALT or telomerase without further details of regulation activity. Thus, the annotation of a given factor in *TelNet* informs if it is involved in ALT or a telomerase dependent TM and how it affects this process. Furthermore, the corresponding *TelNet* score provides an assessment of the significance of this assignment.

**Figure 2.**
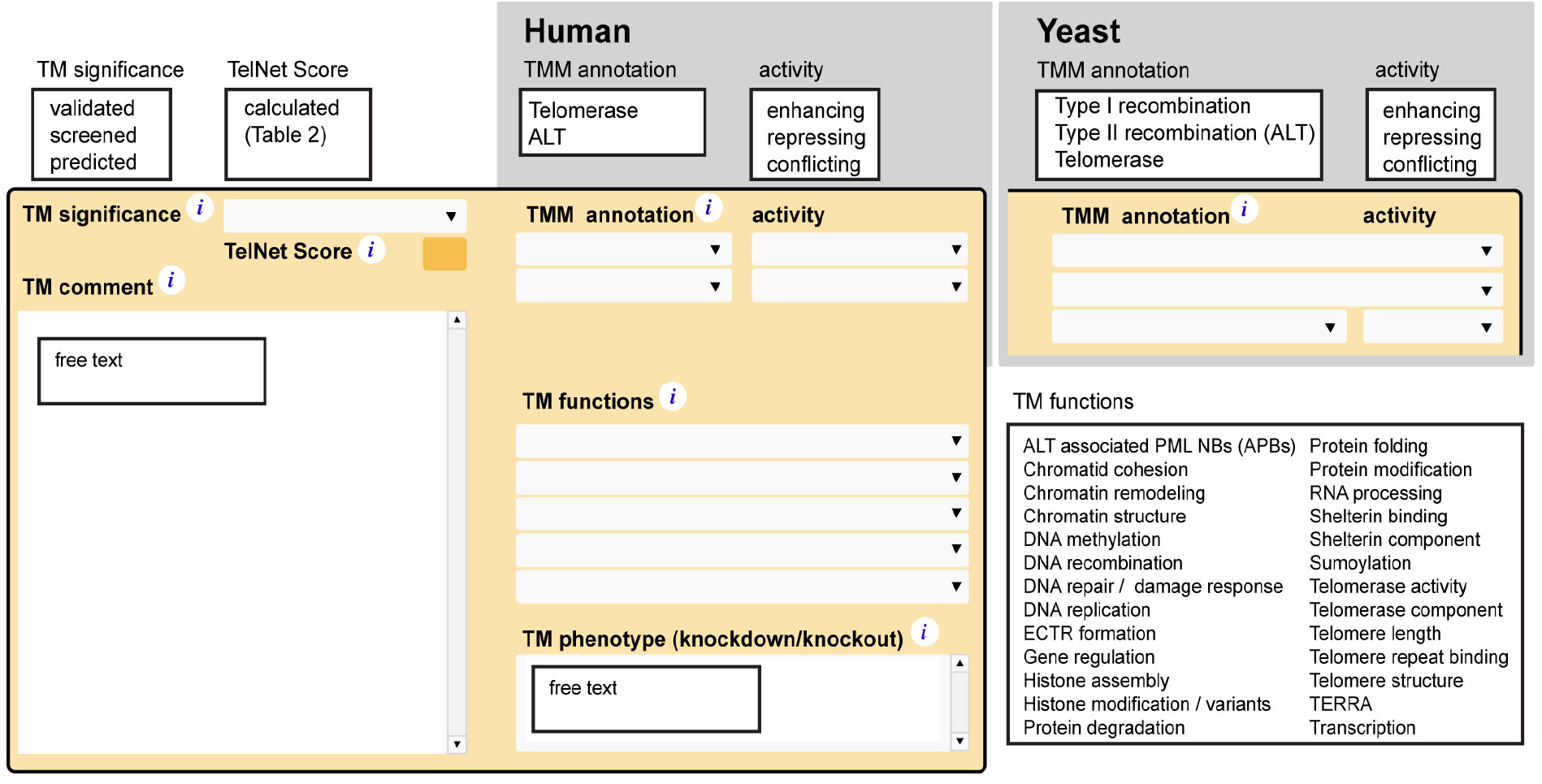
Gene card view of the telomere maintenance information. Annotation fields and possible entries for TM significance, associated *TelNet* score, TM comment, TMM annotation, TM functions and TM phenotype.

**Table 2.**
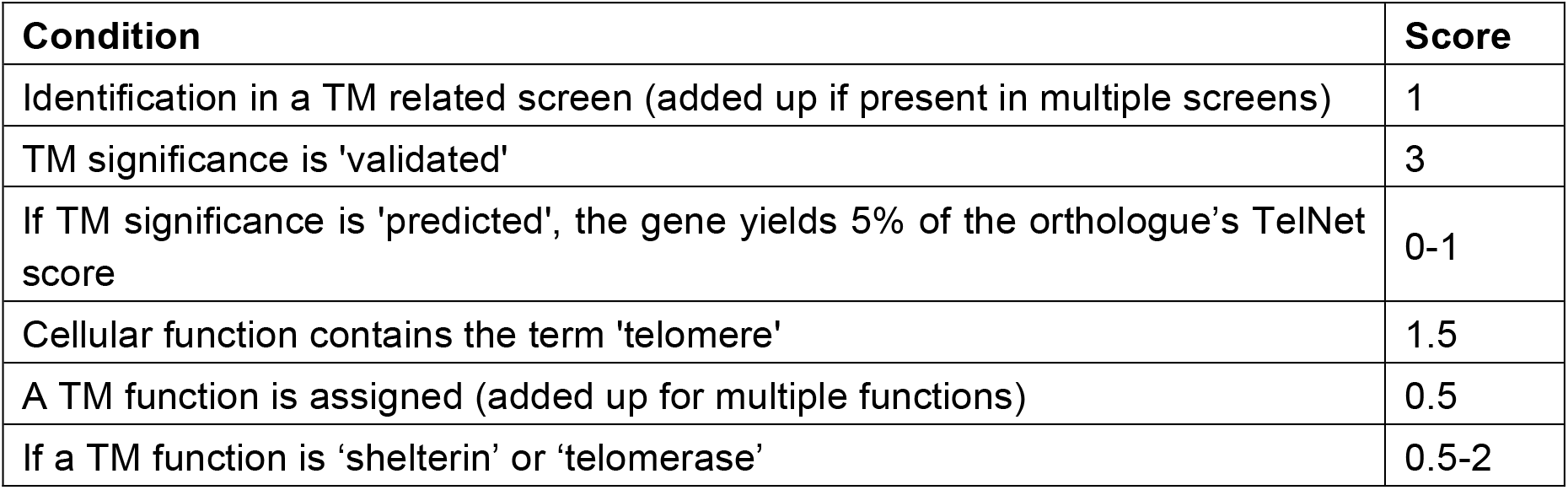
Calculation of the *TelNet* score. The different criteria of evaluated and associated *points* are shown. The *TelNet* score is calculated as the sum of the different entries and can reach a maximum value of 10 points.

## Utility and discussion

### TelNet user interface

On the front layout of *TelNet,* the user must select either *H. sapiens* or *S. cerevisiae* (Figure 3). The default selection is *H. sapiens*. All genes can be browsed by clicking on the “Show all” button. The navigation on top allows switching between different views and returning to the front search page. Gene sets can be displayed as a scrollable list and the complete information of an individual gene is given by selecting the “Card view”. A short explanation of each annotation field is given by clicking on the corresponding info button. Orthologous genes are connected via database hyperlinks. Furthermore, every gene is linked to selected publications.

**Figure 3.**
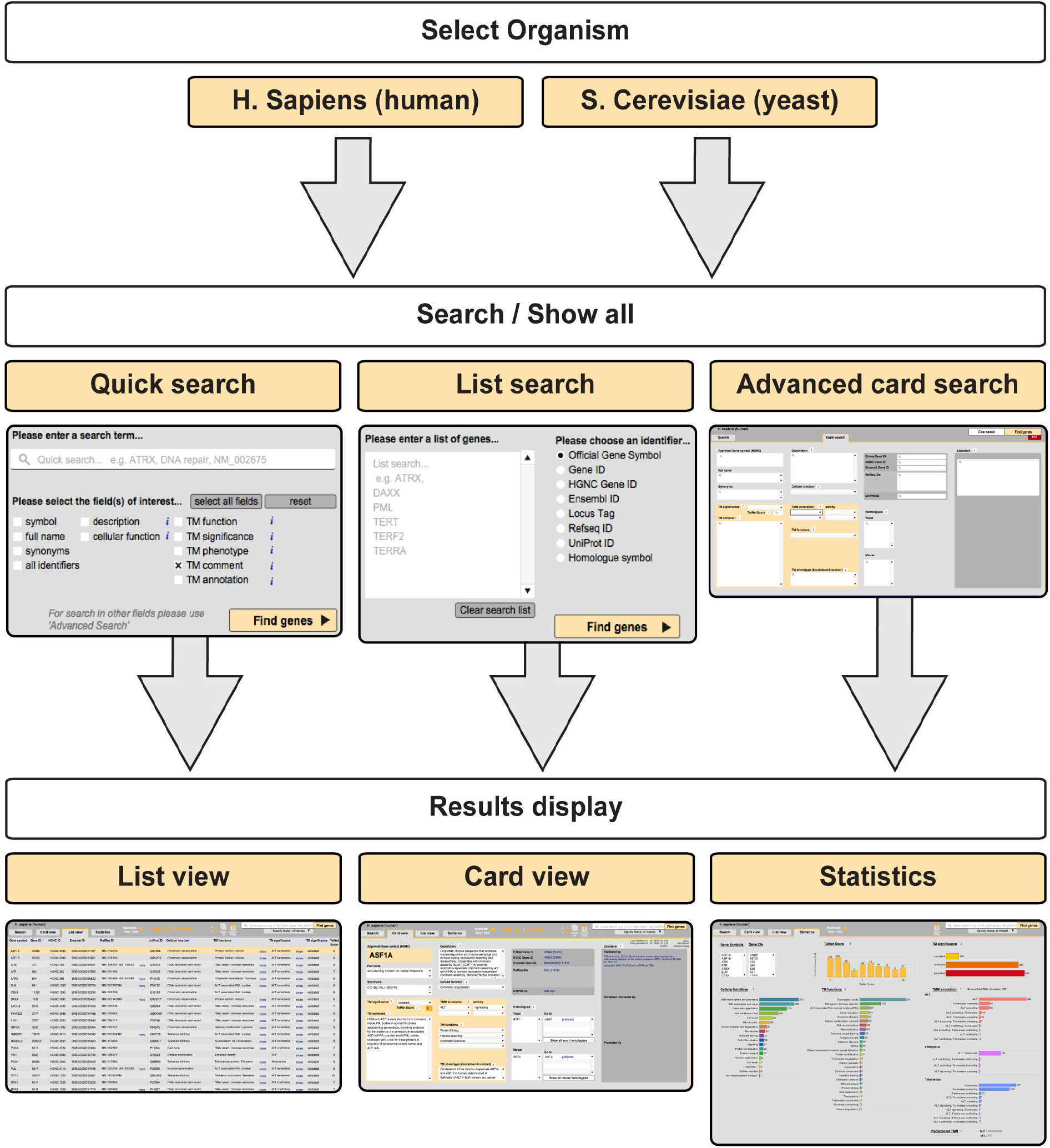
Typical *TelNet* workflow. *Top:* On the front page, the organism is selected. *Middle:* Three different kinds of search tools, namely “quick search”, “list search” and “advanced card search”, are available to retrieve a set of genes. *Bottom:* The resulting genes can be displayed as a scrollable list or as a series of single gene cards. An overview of the associated TM annotations is obtained from the statistics page.

### Search and statistics

The *TelNet* database can be used with three different search modes (Figure 3) named “Quick search”, “List search” and “Advanced card search”. For a quick search throughout selected fields a keyword can be entered into the search bar. If a user wishes to constrain the found set to e.g. a gene name, the selection of fields can be adapted. By performing an advanced card search, the user can enter different search terms in corresponding fields. Furthermore, a complete list of gene identifiers can also be passed into the list search. The organism and identifier provided are mandatory to perform a list search. Genes found are then listed and can be selected for further analysis and TM network identification within *TelNet* or exported in various file formats. The statistics page gives a graphical overview over the distributions of various *TelNet* annotations such as a histogram of the *TelNet* score, and the distribution of TM significance. Furthermore, *TelNet* statistics can be employed for a more detailed pathway analysis regarding TM functions. A predicted wild-type TMM is computed by evaluation of TMM annotations. The wild-type phenotype of a given gene is used for predicting the likely active TMM for a set of genes. Every protein contributes with its *TelNet* score to one of the groups “ALT”, “telomerase-positive” or “ambiguous”, which refers to its wild-type form. Thus, a gene that is recurrently mutated in ALT positive tumors like ATRX (alpha thalassemia/mental retardation syndrome X-linked protein) would represent an ALT suppressor. It is classified as “telomerase-positive” for the predicted TMM associated with its wild-type phenotype. The attribute “ambiguous” is used for genes lacking information on TM mechanism as well as genes with conflicting associations. Thus, *TelNet* informs about known and predicted TM features for the genes of interest via its different search and summary analysis tools.

### Application of *TelNet* for telomere maintenance analysis

In order to illustrate an application of *TelNet*, we performed a pan-cancer correlation analysis of gene expression data with telomere length estimates. It is based on the study of Barthel et al. [39] and uses telomere length data calculated from whole genome sequencing (WGS) data from this reference and gene expression data (stdata_2016_01_28, file uncv2.mRNAseq_RSEM_normalized_log2) downloaded via the Firehose data repository (https://gdac.broadinstitute.org/). A reduced patient data set (*n* = 281) was compiled that comprised samples where normal control samples of matching tumor tissue were available. In order to normalize for tissue- and age-specific effects, we calculated the ratio of tumor over normal for telomere lengths and the log2 ratio for gene expression. For the two ratios the Spearman correlation coefficient was calculated. A significant correlation of telomere length with gene expression was observed for 87 genes (Rho > 0.186 or Rho < −0.184) and genes were differentially expressed (log2 ratio > 0.852 or log2 ratio < −0.782) (Figure 4). It is noted that most of the tumor samples had shorter telomeres than the respective normal control sample. This could be the result of a higher tumor proliferation rate being only partly compensated by the active TMM. This confounding factor as well as the tissue specific expression programs in the different tumor entities are likely to lead to false negative results. For example, TERT (telomerase reverse transcriptase) expression did not show a significant correlation with telomere length. Thus, it might be informative to examine deregulated genes that did not display an (anti-)correlation with telomere length with respect to potential TM activities (**Supplementary Table 1)**.

**Figure 4.**
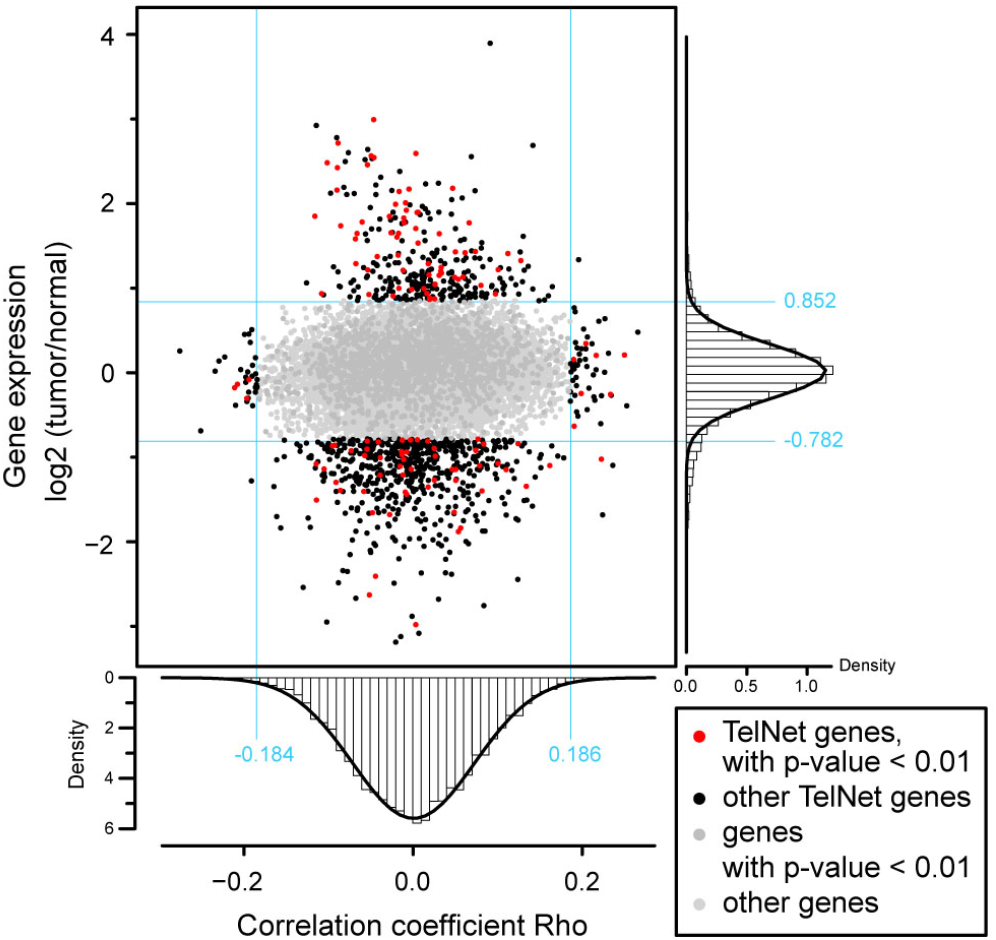
Application of *TelNet* to correlation analysis of telomere length and gene expression. Scatter plot showing the log2 ratio (tumor/normal) of gene expression versus the Spearman correlation coefficient Rho for gene expression and telomere length. For histograms of Rho and log2 ratio a Gaussian fit is shown with significance values defined from the 1%-tail of the fit. Genes that were either significantly (p < 0.01) up- (log2 ratio > 0.852) or downregulated (log2 ratio < −0.782) or significantly (p < 0.01) correlating (Rho > 0.186) or anti-correlating (Rho < −0.184) were colored in black. Genes above the significance thresholds that were present in the *TelNet* database are shown in red color.

To further analyze the genes were correlations were detected, we then used *TelNet* to identify TM links. The *TelNet* “List search” option (Figure 3) retrieved a set of 132 genes (12.9 %). The following results were obtained: (i) None of the genes with a value of the correlation coefficient above the significance thresholds (Rho > 0.186 or Rho < −0.184) was found to be significantly up- (log2 ratio > 0.852) or downregulated (log2 ratio > 0.782) in tumor samples over normal controls. (ii) ARL4D (ADP ribosylation factor-like GTPase 4D) was downregulated in tumors (log2 ratio = −1.02) and expression levels were positively correlated with telomere elongation (Rho = 0.22). Support for the association of ARL4D with telomere length is found by examining the *TelNet* link to the yeast orthologue ARF1 (ADP ribosylation factor 1), which shows shorter telomeres in the deletion mutant [27, 41]. However, the human tumor phenotype appears to be more complex since a downregulation of ARL4D was found in the comparison with healthy tissue. (iii) It appears that from the available pan-cancer data no strong candidate genes are emerging that act as drivers for enhancing or repressing telomere elongation across different tumor-types. Nevertheless, genes like SUMO3 (small ubiquitin-like modifier 3) and ERCC5 (excision repair 5 endonuclease) were identified via the *TelNet* analysis as proteins with known TM activities and are among the most highly (anti-)correlated genes with respect to a function in modulating telomere length in tumors (Table 3). SUMO3 is attached to key proteins of the ALT pathway and shows a positive correlation with telomere length. Sumoylation of PML and shelterin compounds is essential for the formation of PML nuclear bodies [23, 56]. The ERCC5 endonuclease is involved in DNA recombination and repair by annealing single-stranded DNA. Furthermore, ERCC5 positively interacts with the Werner syndrome helicase (WRN) [57] that is involved in telomeric D-loop digestion in ALT cells [58]. Accordingly, a further investigation of SUMO3 and ERCC5 with respect to their role for ALT in tumor cells might be warranted.

**Table 3.**
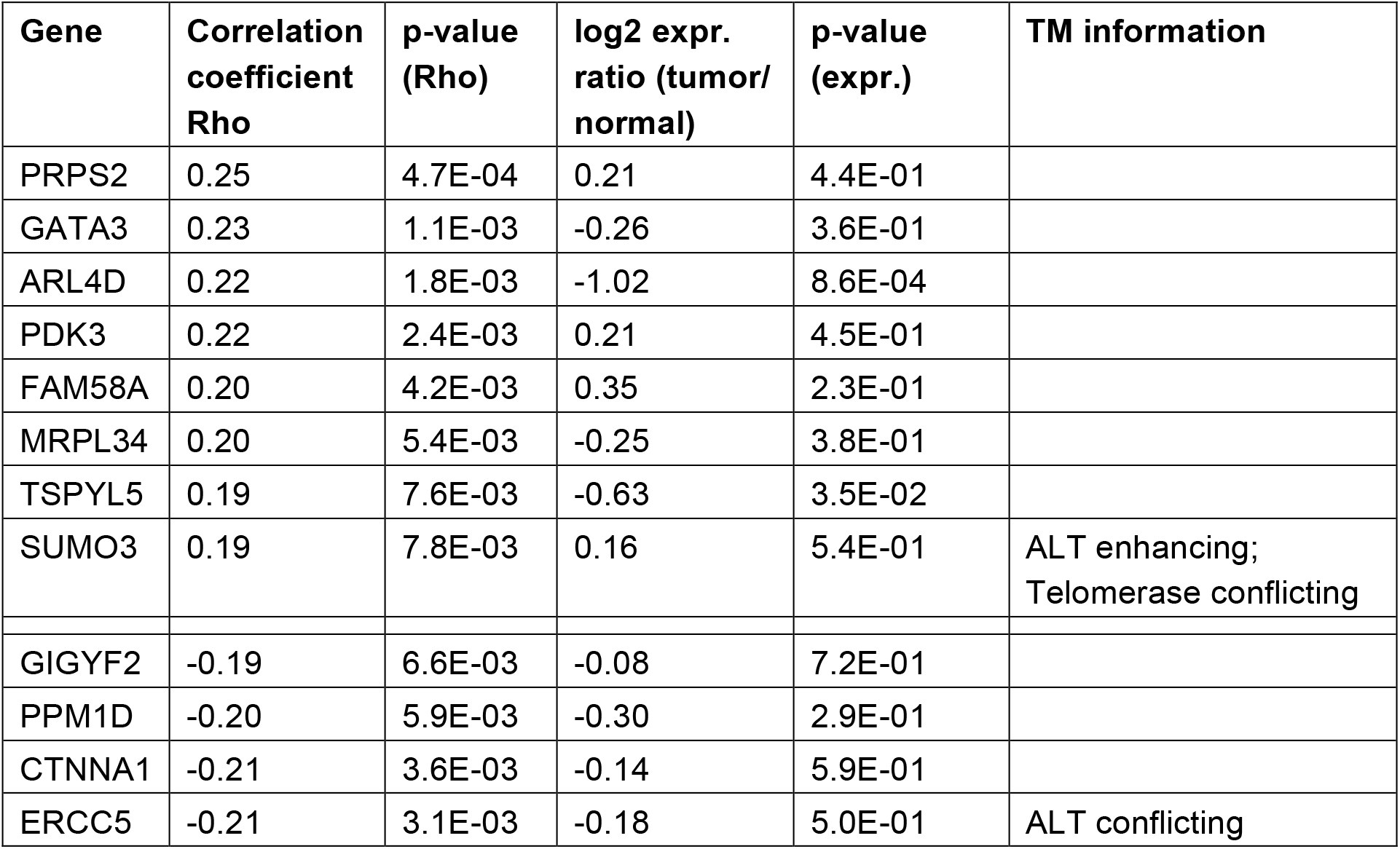
*TelNet* genes (anti-)correlating with telomere length (p < 0.01) changes without altered expression. Significantly (anti-)correlated genes, that are included in the *TelNet* database are shown. Correlation coefficients Rho and the log2 ratio of tumor over normal gene expression are displayed with the respective p-values. Telomere maintenance information from the TMM annotation fields are shown, if available.

In summary, *TelNet* provides a straightforward approach to the annotation of gene lists obtained from different ‘omics’ approaches as illustrated above for WGS and RNA-seq data to identify candidates involved in telomere maintenance for further analysis.

### Future development

The *TelNet* database was developed to provide a comprehensive collection of TM relevant genes. As it is an ongoing research project, new information on telomere maintenance will be added as they become available. In addition, an extension of *TelNet* is planned to include also genes from other organisms such as *M. musculus*. We encourage other researchers working on telomeres to communicate their feedback via e-mail to to further improve the information provided by our database.

## Conclusions

The *TelNet* database compiles information on genes associated to telomeres and telomere maintenance. The annotations provided by *TelNet* allow the distinction between different types of TM mechanisms for a gene (set) of interest according to functional terms and a significance ranking. With these features, *TelNet* supports the identification of TM networks in various ways. A gene set derived from a preceding bioinformatics analysis pipeline can be used for a *TelNet* list search to get more detailed insight on the corresponding TM associated genes. Possible TM links can be explored in an iterative manner. For example, current pan-cancer studies provide a wealth of information on mutated or deregulated genes that can be evaluated in terms of the associated TM mechanisms [39]. Some associations like mutations in *ATRX* and *DAXX* (death-domain associated protein) for ALT as well as *TERT* promoter mutations for telomerase-positive cells are well established. However, in many tumor samples corresponding mutations are absent. One would expect that for these cases the mutation status of a given cancer sample and its active TM are linked via other genes that likely involves a combination of various factors. We anticipate that *TelNet* will prove to be a helpful analysis tool for revealing this type of correlations and to support the identification of active TM networks in different tumor entities.

## Availability of data and material

Database homepage: http://www.cancertelsys.org/telnet/. These data are freely available without restrictions for use in academic research.

### Abbreviations

ALT: alternative lengthening of telomeres
APB: ALT-associated PML nuclear bodies
TM: telomere maintenance
ARL4D: ADP ribosylation factor-like GTPase 4D
ATRX: alpha thalassemia/mental retardation syndrome X-linked protein
ERCC5: excision repair 5 endonuclease
PML: promyelocytic leukemia
TERT: telomerase reverse transcriptase
SUMO3: small ubiquitin-like modifier 3
TM: telomere maintenance
TMM: telomere maintenance mechanism

## Acknowledgements

The authors thank all present and former members of the CancerTelSys Consortium especially Brian Luke, André Maicher, Alexandra Poos, and Rainer König for discussion and data contribution. The authors are grateful to Floris Barthel for providing telomere length data and R scripts for performing the correlation analysis.

## Authors' contributions

DB developed the database and manually included and curated all entries with input from KR, IC and KID. NK was involved in converting the database to the final webversion. DB and KR wrote the manuscript with input from IC. All authors read and approved the final manuscript.

## Consent for publication

Not applicable.

## Ethics approval and consent to participate

Not applicable.

